# Movie-watching evokes ripple-like activity within events and at event boundaries

**DOI:** 10.1101/2023.09.27.559739

**Authors:** Marta Silva, Xiongbo Wu, Marc Sabio, Estefanía Conde-Blanco, Pedro Roldan, Antonio Donaire, Mar Carreño, Nikolai Axmacher, Christopher Baldassano, Lluís Fuentemilla

## Abstract

Ripples are fast oscillatory events widely recognized as crucial markers for memory consolidation and neural plasticity. These transient bursts of activity are thought to coordinate information transfer between the hippocampus and neocortical areas, providing a temporal framework that supports the stabilization and integration of new memories. However, their role in human memory encoding during naturalistic scenarios remains unexplored. Here, we recorded intracranial electrophysiological data from ten epilepsy patients watching a movie. Ripples were analyzed in the hippocampus and in neocortical regions (i.e., temporal and frontal cortex). Our results revealed a differential dynamical pattern of ripple occurrence during encoding. Enhanced hippocampal ripple recruitment was observed at event boundaries, reflecting hippocampal involvement in event segmentation, whereas higher ripple rates were seen within an event for cortical electrodes with higher ripple occurrence at the temporal cortex reflecting weather an event was later recalled. These findings shed light on the neural mechanisms underlying memory encoding and provide insights into the potential role of ripples in the encoding of an event, suggesting an impact in the formation of long-term memories of distinct episodes.

## Introduction

In the process of encoding an event, effective communication between the hippocampus and cortical areas is of utmost importance for memory formation (Baldassano et al. 2017; Geerligs et al. 2021; Ranganath et al. 2005; Reagh and Ranganath 2023). Among the different patterns of neural activity, sharp wave ripples (SWRs) have emerged as a distinct neural signature involved in the transmission of information within the brain. SWRs are characterized by sharp, high-frequency neural oscillations that occur in a highly coordinated and precisely timed manner (Bragin et al. 1999) and can be observed in the local field potential (LFP) signal. Extensive studies conducted in rats have demonstrated that these transient events are accompanied by widespread changes in neural states in both cortical and subcortical regions (Karimi Abadchi et al. 2020; Gomperts, Kloosterman, and Wilson 2015). It is believed that ripples indicate the replay of hippocampal activity and information transfer between the hippocampus and neocortex, facilitating efficient interaction during memory formation and consolidation (Todorova and Zugaro 2020; Vaz et al. 2019; Norman et al. 2021; Dickey et al. 2022). Recent studies have provided evidence for the occurrence of this specific type of neural activity in the human hippocampus (Norman et al. 2019; Vaz et al. 2020; Axmacher, Elger, and Fell 2008), suggesting that it also plays a role in facilitating the formation and retrieval of episodic memories in humans (Sakon and Kahana 2022; Kunz et al. 2024; Sakon et al. 2024). However, whether the formation of memory during naturalistic encoding, where episodic information unfolds continuously, is mediated by ripple events remains unexplored.

Understanding the cognitive and neural underpinnings of episodic memory formation in realistic environments is largely influenced by the view that continuous experiences are rapidly transformed into discrete episodic units via the detection of event boundaries (Zacks et al. 2007). Indeed, event segmentation affects not only our perception of an experience but its subsequent organization in long-term memory (Kurby and Zacks 2008; Radvansky 2012; Sargent et al. 2013), such that elements within an event are bound together more cohesively than elements across events (Ezzyat and Davachi 2011; DuBrow and Davachi 2013; 2014; Horner et al. 2016). Processing at event boundaries has been associated with improved long-term memory for the corresponding event (Newtson and Engquist 1976; Schwan, Garsoffky, and Hesse 2000; Schwan and Garsoffky 2004). Intriguingly, while the hippocampus is particularly active during these moments (Ben-Yakov, Eshel, and Dudai 2013; Baldassano et al. 2017; Ben-Yakov and Henson 2018), neocortical regions have been shown to support memory encoding within events themselves (Reagh and Ranganath 2023). Simultaneously, converging evidence from electrophysiological (Sols et al., 2017; Silva et al., 2019) and functional neuroimaging data (Hahamy, Dubossarsky, and Behrens 2023) has revealed that neural reactivation at event boundaries may be a plausible neural mechanism supporting event memory formation during encoding. Memory reactivation at event boundaries has been shown to engage the hippocampus and neocortical regions (Hahamy, Dubossarsky, and Behrens 2023), regions shown to be involved in ripple-coordinated activity patterns in human (Higgins et al. 2021; Norman et al. 2021), primate (Kaplan et al. 2016), and rodent literature (Karimi Abadchi et al. 2020; Pedrosa et al. 2022). While the underlying functional significance of event boundary-triggered neural reactivation remains uncertain, it has been shown to parallel similar principles accounted for in ripple activity in rodents. Most studies on ripple-based replay have focused on offline periods, such as rest or sleep (Foster 2017; Findlay, Tononi, and Cirelli 2020). However, ripples can also occur in response to significant behavioral occurrences, such as completing a trial, measured as the stopping moment after running from one end of a track to the other (Foster and Wilson 2006), or receiving a reward, after animals traversed spatial trajectories (Singer and Frank 2009). These instances might be akin to event boundaries, delineating contextual shifts in the ongoing experience and reflecting moments where just-encoded information needs to be quickly recapitulated in order to both consolidate it into memory but also to adjust behavior to the new event.

Here, we aimed to examine the dynamical patterns of human ripple occurrence that could support the formation of memories for realistic encoding. Previous research has focused on the detection of ripples during sleep or during awake periods where participants had to encode discrete stimuli. In this study, we investigate, for the first time, the occurrence of this type of neural activity within a continuous and dynamic stream of information. To address this question, we recorded intracranial electrophysiological data simultaneously from the hippocampus, frontal cortex, and temporal cortex of patients undergoing treatment for pharmacologically intractable epilepsy, while they were watching the first 50 min of the first episode of *BBC’s Sherlock*. Building upon earlier evidence of widespread ripple occurrence across the cortex, we examined whether ripple-like activity occurred in diverse cortical areas throughout the encoding of an event and investigated their impact on activity in other regions. Simultaneously, to assess if ripple-like activity reflected the specific hippocampal recruitment at event boundaries seen in previous fMRI studies (Ben-Yakov, Eshel, and Dudai 2013; Baldassano et al. 2017; Ben-Yakov and Henson 2018), we studied how the ripple rate fluctuated around boundaries at the hippocampus and neocortical regions and compared it with the ripple rate within events. Overall, this study provides evidence for a dynamic pattern of ripple-like activity during the encoding of continuous and naturalistic stimuli.

## Results

### Hippocampal and neocortical ripples during movie encoding

To investigate the timing and functional role of ripples during the encoding of naturalistic stimuli, we recorded electrophysiological activity from intracranial electrodes implanted in ten epileptic patients while they watched the first 50 min of the first episode of *BBC’s Sherlock* (**Fig. 1a**), a stimulus already used in previous research (Chen et al. 2017; Baldassano et al. 2017; Silva, Baldassano, and Fuentemilla 2019). They were then asked to freely recall the episode while being recorded using an audio recorder. An event model composed of 38 events and validated in Silva, Baldassano, and Fuentemilla 2019 was used for the current analysis. On average, we found that most of the participants were successful in recalling the encoded events (M = 40.79%, SD = 11.18%), and were accurate in maintaining the order in which the events were presented in the movie during recall (mean Kendall τ= 0.74, p < 0.01), similar to previous findings in healthy participants (Silva, Baldassano, and Fuentemilla 2019).

**Figure 1.**
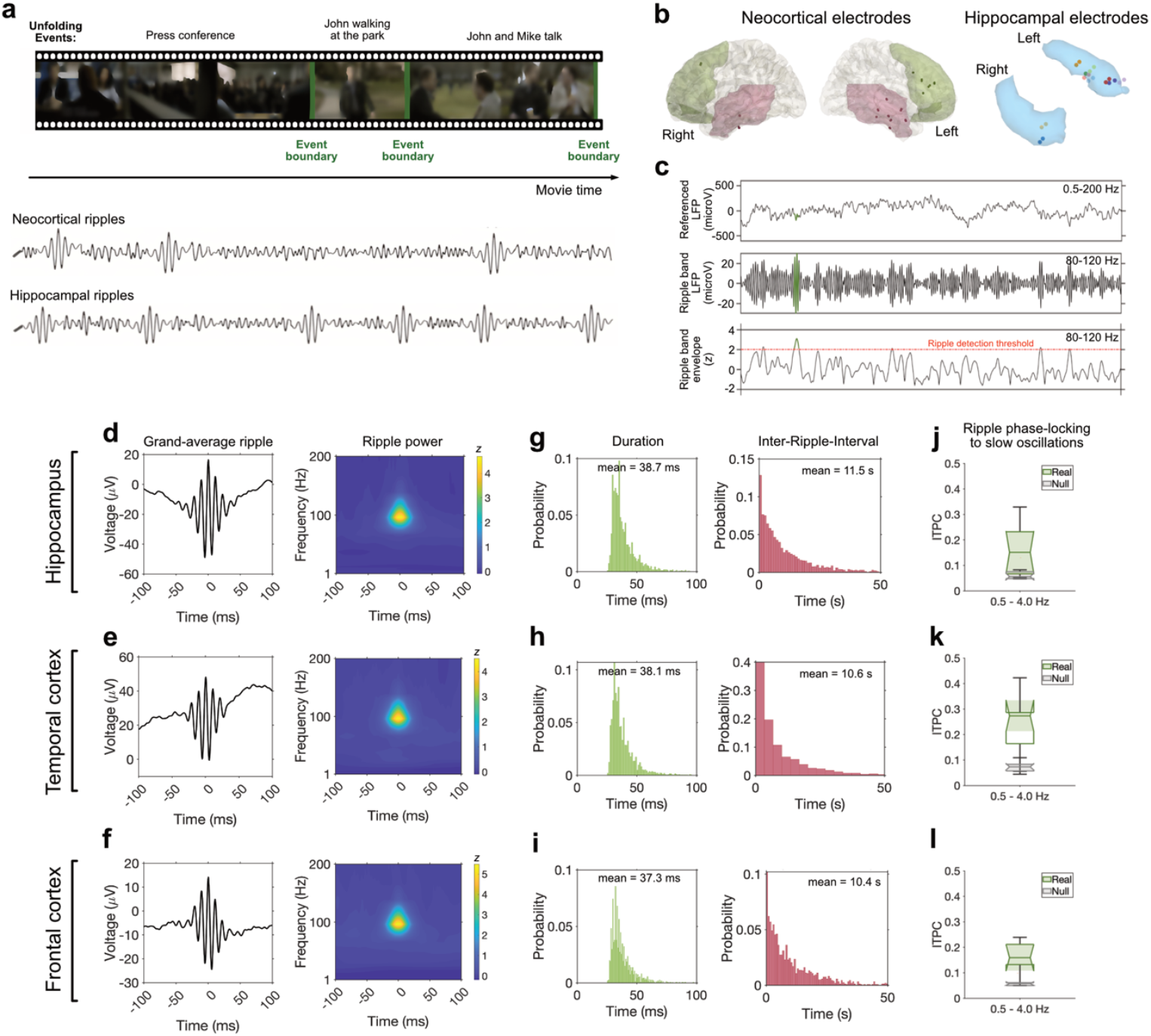
Experimental design, ripple detection and properties. **a)** Experimental design and schematic depiction of expected ripple behavior, with hippocampal ripples occurring closer to event boundaries and neocortical ripples within events. **b)** Left: Temporal cortex (red) and frontal cortex (green) electrode localizations from all participants mapped into common space, shown on a three-dimensional model. Right: Hippocampal electrode locations, each color representing one participant. Each pair of dots indicates the two electrodes from the participants used for bipolar referencing. **c)** Procedure for identifying ripples. Top to bottom: raw LFP; LFP filtered in the 80–120 Hz ripple band; envelope of the ripple-band LFP. In green an example of an identified ripple is shown. **d)** Grand-average voltage trace of hippocampal ripples across all channels in the LFP (<200Hz) time domain and z-scored power spectrogram in the time-frequency domain, with time 0 corresponding to ripple peak. **e)** Similar for temporal cortex and **f)** frontal cortex. **g)** Distribution of ripple durations (green) and inter-ripple intervals (IRIs) (red), across all participants for hippocampus, **h)** similarly for temporal cortex and **i)** frontal cortex. **j)** Inter-trial phase coherence (ITPC) values across ripples (green) and surrogate data (gray), for hippocampus, **k)** similarly for temporal cortex and **l)** frontal cortex. For all boxplots, the central mark is the median, and the edges of the box are the 25th and 75th percentiles.

We identified human ripples during the encoding of the movie by examining LFPs from bipolar macroelectrode channels located at the anterior and middle hippocampus, in either CA1 or adjacent subfields (**Supplementary Fig. 1**), and middle and superior temporal cortex of all ten participants and from electrodes located at rostral medial frontal cortex in six participants (**Fig. 1b**). Following previous ripple detection methods (Vaz et al. 2019; 2020; Kunz et al. 2024) we identified a total of 2444 hippocampal and 2645 temporal cortex ripples across all 10 participants and 1613 frontal cortex ripples across the six participants. The identified ripples exhibited a power peak at ~90 Hz (**Fig. 1d-f, Supplementary Fig. 3**), a mean duration of 38 ms (**Fig. 1g,h,i**), and ripple rates around 5 events/min (Hippocampus = 5.20 events/min; Temporal Cortex = 5.65 events/min; Frontal Cortex = 5.71 events/min) consistent with previous studies in humans (Axmacher, Elger, and Fell 2008; Kunz et al. 2024; Staresina et al. 2015). To confirm that the events used in this study exhibited the narrowband oscillatory behavior characteristic of ripples, we compared the identified ripple candidates to broadband high-frequency events (50–180 Hz). These broadband events were detected using the same algorithm as for ripple detection but with a lower detection threshold (1 SD). By computing a frequency spreading measure we found that the ripples used in the present analysis showed a significantly narrower band than the identified broadband gamma events (**Supplementary Fig. 4)**. The observed Inter-Ripple-Interval (IRI), the time between successive ripples (**Fig. 1g-i**), was ~10 sec and thus slightly longer than reported in previous human studies on task-induced ripple activity (Vaz et al. 2020; Norman et al. 2019; Kunz et al. 2024). This discrepancy suggests that the temporal dynamics of ripples may vary between naturally occurring memory processes and memory formation during cue-locked task conditions.

We next analyzed whether our ripple event candidates occurred predominantly during specific phases of slow band activity (0.5-4Hz), in line with the possibility that ripple occurrence tends to cluster around specific neural states of the ongoing activity (Logothetis et al. 2012a). Consistent with previous findings (Kunz et al. 2024; Mishra et al. 2024), we found that hippocampus and neocortical ripples were phase coupled to ongoing slow oscillations (**Fig. 1j,k,l**), (p < 0.001 for all regions). Ripples in hippocampus and temporal cortex were found to occur at different phases of the ongoing slow oscillations (Kuiper test, p < 0.001; Hippocampal Mean = ~187º; Temporal Cortex Mean = ~311º; **Supplementary Fig. 5b**). In the frontal cortex, ripples aligned with specific phases at individual level as well (Rayleigh test, p < 0.05 for 5 out of the 6 participants; Frontal Cortex Mean = ~140º; **Supplementary Fig. 5b**). Interestingly, however, among the 6 participants, ripples were sometimes locked to the same phases as hippocampal ripples, while they were locked to similar phases as ripples in the temporal cortex in other participants (**Supplementary Fig. 5a**), perhaps due to the inherent variability of the orientation of the implanted electrode in frontal regions. To confirm that the hippocampal ripple phase-locking was specific to a slow band activity, we also assessed to fast theta (4-8Hz) using Rayleigh’s test. We found that ripples were only coupled to slower oscillations (Z=16.99, p<0.001) and not to fast theta (Z=0.24, p=0.78), as in previous sleep studies of ripples (Logothetis et al. 2012b; Sirota et al. 2003; Axmacher, Elger, and Fell 2008; Staresina et al. 2023). Repeating the same analysis but with lower-frequency gamma events (25-75Hz), identified using the same ripple detection method, a phase coupling was present to both slow (Z=25.24, p<0.001) and fast theta (Z=6.33, p=0.002), in line with research showing that high frequency activity of different peak frequencies is phase-locked to the theta cycles (Schomburg et al. 2014). These results provide additional confirmation that our analysis approach might be successfully tracking ripples separately from broadband gamma activity.

Next, we sought to explore whether the frequency occurrence of ripples during encoding of the movie was predictive of memory recollection of movie events during the subsequent recall test. To address this issue, we compared the rate of ripples, normalized by the length of each event, occurring within events that were later remembered or forgotten. Our findings suggest that during the encoding of naturalistic and continuous stimuli, the rate of ripple activity may have a selective direct mechanistic conduit of memory formation depending on the brain region (**Fig. 2**). Ripple occurrence at temporal cortex was significantly higher during the encoding of subsequently remembered versus forgotten movie events (t(9) = 3.367, p = 0.042), whereas no significant differences were found for hippocampal (t(9) = −0.892, p = 0.395) or frontal cortex events (t(5) = 0.566, p = 0.596). To provide a direct test of the interaction between Memory and Region we constructed 4 different linear mixed models (LMMs; **Supplementary Table 2**) that varied in the inclusion of random slopes for each fixed effects (none, Memory only, Region only, both). All models included random intercepts for participants ID. We compared our models by means of the Bayesian Information Criterion (BIC). The model selected (lower BIC) was the one including only one random slope for the effect of Region (BIC = −1688.332, **Supplementary Table 3**). Further, we tested whether the interaction term was significant with a type III analysis of variance. P-values were determined by Satterthwaite’s approximation of degrees of freedom. Results indicated a significant interaction between memory and region (F(2,922.63) = 3.252, p = .039). The current results highlight the relevance of the temporal cortex in determining successful encoding, which is in line with studies in humans showing that the direct stimulation of the temporal cortex, but not the hippocampus or frontal cortex, improves memory formation (Ezzyat et al. 2018; Kahana et al. 2023).

**Figure 2.**
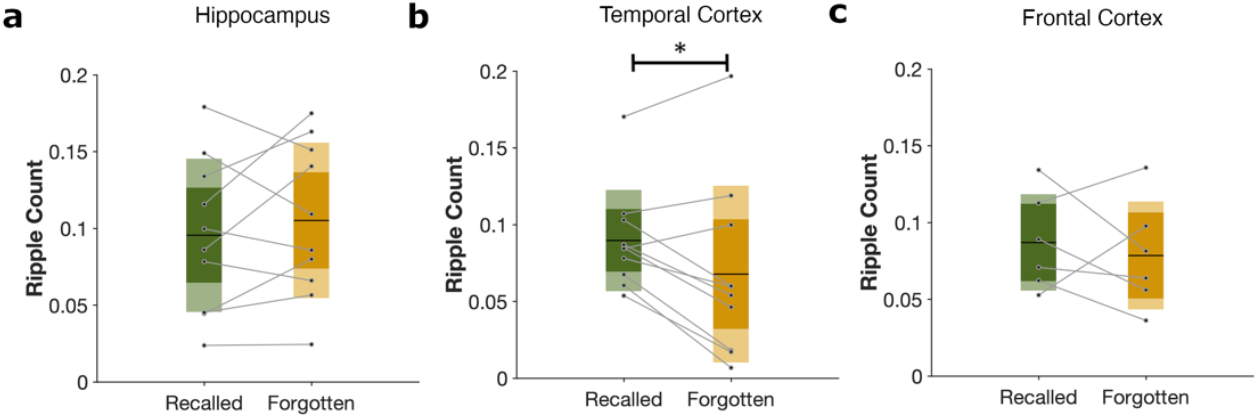
Average frequency of ripples during an event, for each participant, normalized by the length of the event, for recalled (green) and forgotten (yellow) events, in **a)** hippocampus, **b)** temporal cortex and **c)** frontal cortex. For all boxplots, the central mark is the median, and the edges of the box are the 25th and 75th percentiles * Statistically significant at group level (p < 0.05).

### Ripple induced LFP power changes during movie encoding

Having shown that ripples occur during movie encoding in both hippocampus and neocortical regions but exhibit different functional properties, we next examined how they might impact the activity in other regions. Past studies have shown that hippocampal ripples are tightly associated with either the activation or suppression of particular cortical areas (Battaglia, Sutherland, and McNaughton 2004; Sirota et al. 2003; Logothetis et al. 2012a). However, most of these findings are derived from observations of ripple activity during periods of sleep, where ripples align with Slow Oscillations (SO). Given that in our data the ripple candidates show a particular coupling to slow oscillations, even during active processing, similar to previous research (Kunz et al. 2024), we explored how the interaction of hippocampal and neocortical ripples was related to neural state changes in different regions by looking at how the LFP signal from one region might be altered when a ripple is occurring in another region. For that, we computed the time-frequency spectrum in one region time-locked to a ripple occurrence in one other region (e.g. the time-frequency at a window of time in the temporal cortex around a ripple occurring at the hippocampus).

We first examined LFP power changes in cortical regions locked to the occurrence of hippocampal ripples. This analysis revealed a reduction of LFP power in high frequencies (>32Hz) of the temporal cortex around hippocampal ripples (cluster permutation test, tmax = 4.376, tmean = −2.812, p < 0.001, **Fig. 3a**). This decrease in LFP power was not, however, present at frontal regions around hippocampal ripples, where no significant clusters were identified (**Fig. 3b**). These results are consistent with recent rodent studies highlighting that, unlike during sleep, neocortical activity is dominated by inhibition around awake ripples (Karimi Abadchi et al. 2023).

**Figure 3.**
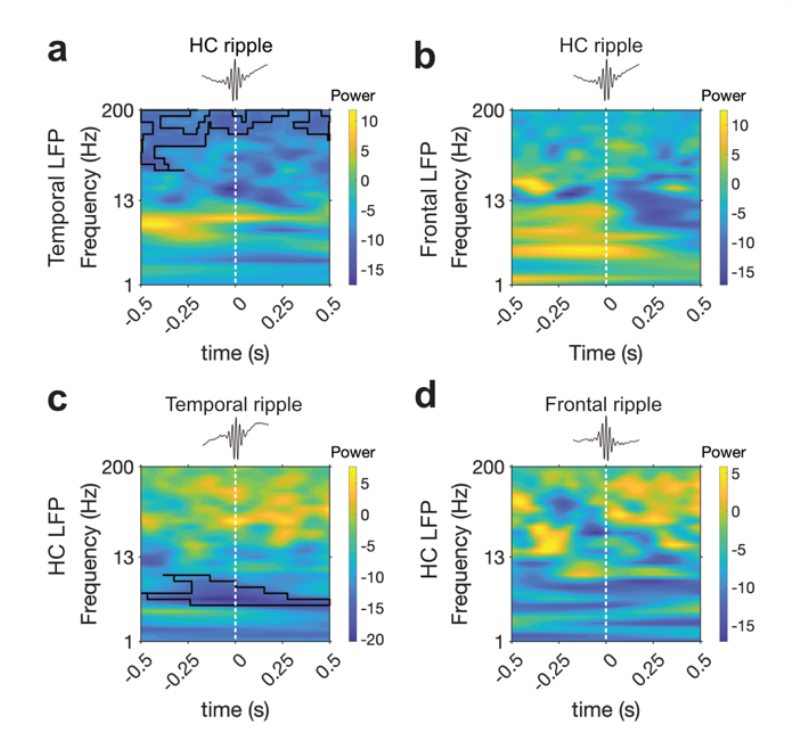
LFP power (z-scored) changes in (**a**) temporal cortex and **(b)** frontal cortex locked to hippocampal ripples, where 0 corresponds to ripple peak. LFP power (z-scored) changes in hippocampus locked to (**c**) temporal cortex and **(d)** frontal cortex ripples during the encoding of the movie, where 0 corresponds to ripple peak. Black contours correspond to statistically significant clusters (two-sided cluster-based permutation tests: p < 0.05). Data in all above analysis was smoothed with a gaussian filter across time (kernel length, 0.2 s).

Having observed that temporal cortex activity is strongly modulated during hippocampal ripples, we investigated whether the reverse was also the case – i.e., whether hippocampal activity was modulated during neocortical ripples. This analysis revealed a marked decrease of hippocampal LFP power at low frequencies corresponding to the alpha band (7-12Hz) around the onset of temporal cortex ripples (cluster permutation test, tmax = 5.345, tmean = −3.342, p = 0.012, **Fig. 3c**). Alpha band oscillatory power decreases have been implicated in memory encoding (Paller and Wagner 2002; Kim 2011) and proposed to act as a gating mechanism whereby decreasing alpha power increases firing rates (Hanslmayr, Staresina, and Bowman 2016). These results propose that temporal cortex ripples might be decreasing alpha power in the hippocampus in order to potentiate memory encoding. A similar time-frequency map is observed during frontal cortex ripples, with higher power for higher frequencies and lower power for low frequencies. However, no significant cluster was found after performing the permutation test (**Fig. 3d**).

We computed interaction tests between LFP changes and regions by computing the difference between the time-frequency maps of the two rows on **Fig. 3** (3a – 3b and 3c – 3d) and performing the same cluster permutation analysis as used previously to look for clusters across frequencies and time. We tested for a difference in spectral power changes between temporal and frontal regions, time-locked to hippocampal ripples, and also tested whether spectral power changes in the hippocampus differed when time-locking to temporal cortex ripples versus frontal cortex ripples (**Supplementary Figure 8**). For both interaction tests we found no significant clusters, raising the possibility that the lack of frontal cortex effects in **Fig. 3b and 3d** could be due to sample size limitations (since only six participants had recordings from frontal cortex) rather than reflecting a meaningful difference between the hippocampus and frontal cortex.

### Hippocampal ripples increase around movie event boundaries

Event boundaries – i.e. time points at which there is a shift in one’s current event model – are thought to be the moments in time when the organization and binding of information into long-term memory occurs (Ben-Yakov and Henson 2018; Baldassano et al. 2017; Silva, Baldassano, and Fuentemilla 2019). In line with findings in rodents that hippocampal ripples promote memory formation for just encoded events (Foster and Wilson 2006; Diba and Buzsáki 2007; Karlsson and Frank 2009), we next tested whether the occurrence of hippocampal ripples increased around event boundaries during movie viewing (Bilkey and Jensen 2021), possibly reflecting brief temporal opportunity windows of memory plasticity during awake encoding (Foster 2017). To test this hypothesis, we calculated a peri-boundary ripple rate by computing the peristimulus time histogram (PSTH) relative to all boundary onsets and compared it to a null distribution by shuffling the temporal order of the events while maintaining their lengths. We found a marked increase in hippocampal ripple activity concomitant with a reduction in neocortical electrode activity specifically at boundaries (**Fig. 4**). The increase of hippocampal ripples at boundaries is in line with fMRI studies which show that stronger event encoding was related to lower hippocampal activation during the event and a high activation at its offset (i.e., at boundaries) (Ben-Yakov and Henson 2018; Baldassano et al. 2017). The decreased activity in the cortex could be indicative of a switching mechanism in which neocortical regions need to be silenced at moments in which resources have to be concentrated in the hippocampus (Logothetis et al. 2012a).

**Figure 4.**
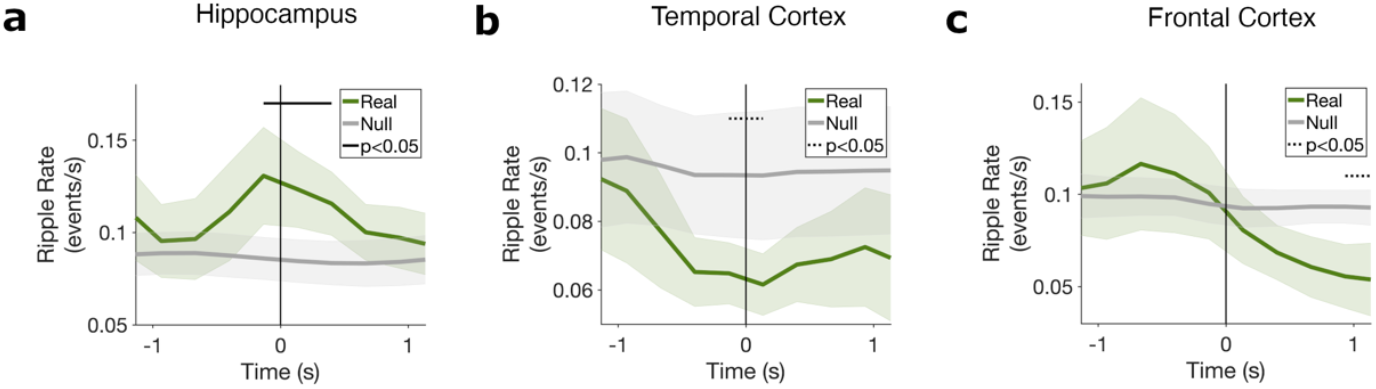
Instantaneous ripple rate computed in 300ms time bins around boundary onset and smoothed by a five-point triangular window, for empirical data (green) and surrogate data (grey), for **a)** hippocampus, **b)** temporal cortex and **c)** frontal cortex. Shaded region corresponds to SEM across participants. Black line at top indicates significant clusters with FDR correction (p < 0.05) and black dashed line at top indicates significant clusters with no FDR correction (p < 0.05).

## Discussion

In this study, we recorded intracranial electrophysiological data from the human brain to examine the properties and interactions of ripple-like events in hippocampus and in frontal and temporal cortex while participants watched a continuous 50 min movie. We found that ripples occurred in both hippocampus and neocortical areas during the encoding of a movie event. Hippocampal ripple activity increased at event boundaries whereas cortical ripples were recruited within an event, reflecting a distinctive rhythm of ripple timing during continuous perception. Additionally, ripple occurrence had a differential impact on the memory encoding of an event, as only ripples occurring in the temporal cortex predicted later recollection of that event. This differential cortico-hippocampal pattern of ripple activity during encoding highlights the involvement of ripples in the formation of episodic memories in naturalistic circumstances.

The hippocampus has been found to exhibit heightened activity and engagement at event boundaries (Ben-Yakov, Eshel, and Dudai 2013; Baldassano et al. 2017; Ben-Yakov and Henson 2018), while other cortical areas appear to be more sensitive to representing information within an event (Reagh and Ranganath 2023). In our study, we observed that this alternating recruitment pattern during the encoding of an event is reflected at the level of ripple activity. Specifically, when examining event boundaries, we observed an increase in hippocampal ripples. Hippocampal ripple probability has been shown to increase in a pre-stimuli window where uncertainty was high (Frank et al. 2023). Event boundaries are often points in time when activity becomes less predictable (Swallow, Zacks, and Abrams 2011). The increase of hippocampal ripples seen at event boundaries might then suggest a role of hippocampal ripples in reporting the predictability of an upcoming stimulus. Event boundaries were also accompanied by a decrease in cortical ripples. The recruitment of ripples at different moments and in different areas during the encoding of a dynamic event may serve as a computationally efficient strategy for simplifying complex events into key components. Furthermore, the coactivation of stimulus-specific cells during hippocampal ripples, as observed during the encoding of object-place associations in humans (Kunz et al. 2024), suggests that ripples may play a facilitative role in binding diverse memory elements represented across distinct cortical areas. This facilitation enables the formation of coherent event representations, supporting the integration of information from different cortical regions. Simultaneously, waking SWR might serve as a neurophysiological mechanism for memory selection by tagging aspects of experience that should be preserved and consolidated for future use (Yang et al. 2024).

The investigation of ripple events during awake behavior in rodents have shown that ripples often occur in instances which might be akin to event boundaries, delineating contextual shifts in the ongoing experience, such as completing a trial (Foster and Wilson 2006) or receiving a reward (Singer and Frank 2009). These periods are often punctuated by structured and temporally-compressed replay of hippocampal multi-cell sequences representing past navigation-related experiences, as well as “preplay” of potential future (Diba and Buzsáki 2007; Pfeiffer and Foster 2013; Jadhav et al. 2012; Foster and Wilson 2006; Gupta et al. 2010). In humans, boundaries have been shown to trigger a rapid reinstatement of the just-encoded event (Sols et al. 2017; Silva, Baldassano, and Fuentemilla 2019; Wu et al. 2022; Hahamy, Dubossarsky, and Behrens 2023). The observed increase in ripples at boundaries may indicate a window of enhanced hippocampal cell recruitment needed for a replay-like mechanism. However, the link between this increase in ripples and either boundary-triggered reinstatement patterns or subsequent memory effects remains unclear. Given the typically low ripple rates seen in humans (rates of ~0.1Hz) and the limited number of event boundaries in this dataset, we could not assess ripple rates separately for recalled and forgotten events, as this would yield an extremely low number of ripple events within a brief window around event boundaries. Further research is needed to determine whether there is a connection between these two phenomena and whether there is any link between ripple occurrence throughout event encoding and replay. Shedding light into this direction may help characterize the functional properties of ripple activity in complex memory processes that may be difficult to assess experimentally in the rodent research field.

Event-specific patterns of brain activity are observed across widespread regions of the brain during naturalistic movie tasks. Studies have suggested that certain components of the default-Mode Network (DMN) – and particularly, the medial prefrontal cortex (mPFC), represent abstract information about events that might generalize across instances of similar situations (Baldassano, Hasson, and Norman 2018; Reagh and Ranganath 2023). Conversely, an anterior temporal (AT) network is thought to represent information about various categories including objects and people (Peelen and Kastner 2011; Tsantani et al. 2019; Reagh and Ranganath 2023). The communication between these brain regions during event encoding ensures the integration of perceptual details with contextual information, facilitating the formation of meaningful and distinct memory traces. One mechanism through which the brain might achieve this integration is through sharp wave-ripples. Ripples are not isolated hippocampal events but are part of a complex network of interconnected oscillatory assemblies involving both neocortex and hippocampus (Dickey et al. 2022). The coordination of these networks facilitates specific information transfer between neocortical and hippocampal cell assemblies, contributing to the coherent processing and encoding of event-related information. Given these observations and our electrode coverage, we decided to analyze data from the anterior and middle temporal cortex and from middle frontal areas to assess how ripple activity might be recruited differently throughout the encoding of an event.

We observed a pattern of ripple activity occurring in the temporal cortex, specifically during events that were later successfully recalled. The temporal cortex, and in particular the anterior temporal lobe, where the majority of electrodes used in this analysis were placed, plays a critical role in semantic memory and represents information about objects and individuals (Bonner and Price 2013; Reagh and Ranganath 2023). The occurrence of ripple activity in the temporal cortex during the encoding of an event might suggest its involvement in capturing this type of information. During the event encoding process, cortical ripples may be triggered by stimulus-specific neuronal activity, facilitating the transfer of information from extrahippocampal regions to the hippocampus.

Similarly, we observed that ripples in the frontal cortex were particularly recruited during the encoding of an event. The frontal cortex receives direct and indirect projections from the hippocampus (Cenquizca and Swanson 2007) and has been implicated in various functions such as decision-making, long-term memory consolidation, and working memory (Cenquizca and Swanson 2007). The majority of frontal electrodes included in this analysis were located in the rostral medial cortex, which have also been associated with prospective memory (Volle et al. 2011). Although the precise role of the frontal cortex in the current task remains uncertain, it seems reasonable to hypothesize its involvement in monitoring working memory maintenance during an event (Kurby and Zacks 2008; Radvansky 2017).

Furthermore, in line with recent research on human ripples (Kunz et al. 2024; Tong et al. 2021; Mishra et al. 2024), we found that ripples occurred in a phase-locked manner to specific phases of slow oscillatory activity. During sleep, ripples generally appear at the descending phase of neocortical delta oscillations (Battaglia, Sutherland, and McNaughton 2004; Staresina et al. 2015; 2023), which have been associated with alternating states of enhanced and reduced cortical excitability (Steriade, Nunez, and Amzica 1993). Alternatively, the frequency band associated with delta (0.5-4Hz) during awake behavior has been shown to reflect slow theta oscillations (1-4Hz) in humans. Particularly, human hippocampal slow 1 - 4 Hz oscillations have been shown to have similar functional properties to the theta oscillations observed in rodents (Jacobs 2014) and to be related to successful memory encoding (Rudoler, Herweg, and Kahana 2023). Theta oscillations have been suggested to underlie active memory function whereas ripple oscillations represent offline memory function. This raises the concern that the effects seen in our analysis are reflecting a hippocampal theta-gamma coupling instead. Gamma patterns at all frequencies are phase-locked to the theta cycles (Schomburg et al. 2014). In contrast, SPW-Rs in sleep seem to be absent during theta but coupled with delta waves (Sirota et al. 2003; Axmacher, Elger, and Fell 2008). Our results show a similar pattern during awake behavior, where slower gamma events seem to be phase-locked to theta cycles, both slow and fast, whereas ripple events are only phase-coupled to slow oscillations. Together with the previous studies showing coupling between ripples and slow oscillations (Kunz et al. 2024; Mishra et al. 2024), our findings point to the possibility that, in the awake the phase-coupling with slow oscillations might have a potential different role than the slower gamma events phase-locking to theta.

We also observed widespread changes in LFP that were linked to the occurrence of ripples. When a ripple occurred in the hippocampus, temporal cortex high-frequency activity was suppressed, and conversely, when a ripple occurred in the temporal cortex, hippocampal activity showed a decrease in alpha power, which has been associated in the past with successful encoding (Paller and Wagner 2002; Kim 2011). Our results show a opposite effect to the one observed in previous studies where power at lower frequencies decreased and power at higher frequencies increased during hippocampus ripples (Kunz et al. 2024). Nevertheless, this discrepancy might simply suggest that the impact of ripples in different cortical areas may vary between naturally occurring memory processes and memory formation during cue-locked task conditions.

Overall, our observations suggest the existence of a dynamic system where the occurrence of ripples may influence the silencing or activation of specific brain areas and impact memory differently. Research indicates that hippocampal SWRs are synchronized with neocortical ripples during memory tasks and influenced by neocortical slow oscillations and spindle frequency oscillations during non-rapid eye movement sleep (NREM) (Staresina et al. 2023; 2015; Dickey et al. 2022). Although it is unclear if the cortical events are precisely the same as hippocampal ripples, these brief bursts of fast oscillations appear to play a functional role. Both cortical and hippocampal ripples have been shown to occur, in both NREM and waking, during sudden increases in local excitability lasting at least as long as the ripple (~70 ms, Dickey et al. 2022) and to synchronize during memory tasks (Vaz et al. 2020; Norman et al. 2021). Waking cortical ripples have also been shown to reflect spatiotemporal firing patterns during the cued recall of items which replicate the patterns initially evoked by those items during their first presentation (Jiang et al. 2017). This suggests that ripples across different conditions—humans and rodents, NREM and waking, hippocampus and cortex—might share a common role in reconstructing previously occurring firing patterns. The similarity in the characteristics of human ripples, whether in NREM or waking states, and whether in the hippocampus or cortex, supports this idea. Cortical ripples have been shown to be similar to hippocampal ripples in oscillation frequency, density, and duration (Dickey et al. 2022). Our study provides further evidence suggesting that these oscillations exhibit similar characteristics in duration and frequency to hippocampal ripples and reflect the activation of these brain regions during event encoding, impacting subsequent recall. However, the constrained coverage of iEEG recordings in humans, dictated by clinical electrode placement, precludes drawing conclusions about ripple activity in other neocortical regions, such as the precuneus or angular gyrus, despite their consistent involvement in event segmentation during task-based movie clip studies (Chen et al. 2017; Baldassano et al. 2017).

There is an ongoing debate questioning whether human “ripples” align with rodent counterparts (Liu et al. 2022). Central to this discussion is the challenge of establishing, in humans, the precise physiological characterization necessary for discerning ripple activity and discerning it from other high-frequency patterns, such as high frequency oscillations (HFO), or epileptic activity. Most of the difficulties encountered in the human research field are due to technical and ethical limitations, as intracranial recordings are strictly guided by clinical purposes, and therefore, electrode localization, size of the electrodes (macro vs micro), and the ability to sample extensive brain regions over prolonged periods is very limited in humans compared to rodent research. Nevertheless, several methodological strategies have been proposed to address this issue (Liu et al. 2022). While valuable, some of them have been difficult to put into practice. For example, surgical implantation in human patients that targets the hippocampus tends to often, almost exclusively, reach the anterior part of the hippocampus, as is the case for the electrodes used in our study. However, the human anterior hippocampus corresponds to the ventral hippocampus in rodents, whereas rodent sharp-wave ripple studies have been mostly recorded from the dorsal hippocampus of rodents. This means that direct comparisons between the ripples identified in human studies and sharp-wave ripples reported in rodent studies have not been possible. Second, the relationship between ripples and HFOs warrants further investigation, as theta-gamma coupling can potentially be an alternative mechanism associated with event boundary memory processes. The events used in this study show narrow-band increases in oscillatory activity (**Supplementary Fig. 4)** within the same band as previous human studies which used the same detection algorithm. Additionally, hippocampal events were only phased-locked to slow oscillations and not to fast theta, as seen previously in sleep (Logothetis et al. 2012a; Sirota et al. 2003; Axmacher, Elger, and Fell 2008; Staresina et al. 2023), and have similar ripple rates to previous sleep studies in humans (Staresina et al. 2023; Axmacher, Elger, and Fell 2008). Nevertheless, in the case that the events used in the study are not truly ripples and instead represent theta-gamma coupling, both mechanisms might be useful for memory encoding of events: theta-gamma could be used to organize the order in which the elements that comprise an event are encoded or to link different events together (Griffiths and Fuentemilla 2020), whereas ripples could serve as a way to integrate new information by tagging which aspects of experience should be preserved and consolidated for future use (Yang et al. 2024). Disentangling the two and testing this complementary role warrants further investigations. Finally, ripples are also dynamic entities showing different morphologies, such as different amplitudes and durations, based on brain region or behavior (Buzsáki 2015). In applying fixed criteria, as in the currently employed detection methods, we might risk overlooking functionally meaningful events. Nevertheless, to validate the ripples identified with the chosen detection method (Vaz et al. 2020; Kunz et al. 2024) we have repeated and validated the analysis reported in this manuscript with a distinct detection method (Norman et al. 2019, **Supplementary Fig. 9)**. Unfortunately, additional recommended methodological strategies, such as a direct comparison between ripples from healthy and diseased hippocampus or comparing ripple properties from NREM and awake states (Liu et al., 2022) were not easily feasible in human research given the strict data availability in clinical settings. Future studies are essential for a deeper understanding of the underlying nature of ripples in the context of naturalistic encoding in humans.

In sum, our results provide the first evidence for the involvement of ripple-like activity in the formation of episodic memories in naturalistic circumstances. We observed increased ripple activity at event boundaries in the hippocampus and within events in cortical regions, reflecting distinctive patterns of information processing during different event periods. Furthermore, the occurrence of ripples had varying effects on the memory encoding of an event, with only ripples in the temporal cortex being indicative of later recollection of that event. These findings shed light on the intricate mechanisms underlying memory encoding and provide insights into the potential role of ripples in event segmentation and memory.

## Data Availability

Due to ethical restrictions from the Ethics Committee, structural data cannot be made publicly available as it contains information related to the privacy of the research participants. Data files for the three regions used in these analyses will be included in a CodeOcean capsule.

## Code Availability

Code for this manuscript’s analyses will be available as a CodeOcean capsule upon publication.

## Acknowledgments

This work was supported by the Spanish Ministerio de Ciencia, Innovación y Universidades, which is part of Agencia Estatal de Investigación (AEI), through the project PID2019-111199GB-I00 to L.F. (Co-funded by European Regional Development Fund. ERDF, a way to build Europe). We thank CERCA Programme/Generalitat de Catalunya for institutional support. The project that gave rise to these results received the support of a fellowship from “la Caixa” Foundation (ID 100010434). The fellowship code is “LCF/BQ/DI19/11730040”.

## Author Contributions

M.Si. and L.F. designed research; M.Si, X.U. and M.Sa collected the data; E.C, P.R., A.D. and M.C implanted electrodes; M.Si., X.U. and L.F. analyzed data; M.Si., N.A., C.B., discussed the results; M.Si. and L.F. wrote the paper. All authors reviewed and revised the final manuscript.

## Competing Interests

The authors declare no competing interests.

## Materials and Methods

### Data collection

We tested 10 human subjects who were undergoing treatment for pharmacologically intractable epilepsy at Hospital Clínic – IDIBAPS in Barcelona. Prior to performing the task, all participants were thoroughly briefed on the specificities of the task, ensuring they had a comprehensive understanding of the objectives, procedures, and potential risks involved. Each participant was provided with a consent form, which they attentively reviewed and signed, demonstrating their informed consent to participate in the study. The study was approved by the hospital Ethics Committee.

Patients were surgically implanted with intracranial depth electrodes for diagnostic purposes to isolate their epileptic seizure focus for potential subsequent surgical resection. Antiseizure medications were reduced to record seizures during the patient’s hospital stay. Cognitive testing was performed ≥7h after the last seizure (Henin et al., 2021). The exact electrode number and locations varied across subjects and were determined solely by clinical needs. The recordings were performed using a clinical EEG system (Natus Quantum LTM Amplifier) with a 2048Hz sampling rate and an online bandpass filter from 0.1Hz to 4000Hz. Intracerebral electrodes (Microdeep, DIXI Medical) were used for recordings. Each multielectrode had 8 to 18 contacts, spaced 5 mm and 1 to 2 mm long with a diameter of 0.8 mm.

### Experimental Design

The experiment was conducted in a sound-attenuated room in the hospital, with participants sitting upright in a comfortable chair or on their bed. Participants were asked to watch the first 50 min of the first episode of *BBC’s Sherlock*, dubbed in Spanish, as done previously in Silva, Baldassano, and Fuentemilla 2019. Participants were informed that a subsequent recall memory test would follow. After the movie, some time was given to rest (5-10 min) before the test began. During the test, they were asked to freely recall the episode without cues while being recorded using an audio recorder placed on the overbed table next to the laptop computer. The audio files were later analyzed to access the participants’ length of the recall. The experimental design was implemented using PsychoPy (Peirce et al. 2019) and presented on a 13-inch portable computer, placed on the overbed table at approximately 60 cm distance in front of the patients.

### Event boundary annotations

The event model validated in Silva, Baldassano, and Fuentemilla 2019 was used for the current analysis. This model is composed by 38 events (minimum = 4 s, maximum = 444 s, and mean = 76.02 s) and it was constructed by having six external participants annotate the temporal point at which they felt “a new scene is starting; these are points in the movie when there is a major change in topic, location or time” (Baldassano et al., 2019). The final model was built based on boundary time points that were consistent across observers. To find a statistical threshold of how many observers should coincide in a given time point to be different from chance in our data, we shuffled the number of observations 1000 times and created a null distribution of the resulting coincident time points. An α = 0.05 as a cutoff for significance indicated that boundary time points at which at least 3 observers coincided in (considering 3 s as possible window of coincidence as in Baldassano et al. 2017) could not be explained by chance.

### Verbal recall analysis

The audio files from the free verbal recall were analyzed by a laboratory member who was a proficient Spanish speaker, using the list of events from the event model mentioned in the previous section. An event was counted as recalled if the participant described any part of that scene.

To statistically assess whether the order of events during movie watching was preserved during free recall, we computed Kendall rank correlation coefficients between each individual event temporal order and a simulated correct linear order. A positive Kendall tau coefficient close to 1 indicates that the encoded temporal order of the events was highly preserved during their recall.

### Electrode localization and selection

The presence of electrodes in the respective brain areas was assessed with the examination of a computed tomography (CT) and preoperative Magnetic Resonance Imaging (MRI) T1 scans. Cerebral atlases of each patient were obtained with the parcellation of the preoperative T1 using Freesurfer (https://surfer.nmr.mgh.harvard.edu). The CT was co-registered to the T1 and contact tags and names were placed manually using fieldtrip’s toolbox (https://www.fieldtriptoolbox.org/). Hippocampal segmentation was performed using Freesurfer’s parcellation function. Selection of channels was done in native space to prevent errors due to distortions.

To eliminate potential system-wide artifacts or noise and to better sense ripples locally, we applied bipolar re-referencing between pairs of neighboring contacts. The channels of interest were selected based on the following criteria: if more than one channel was eligible, we privileged the channel that had an adjacent distal referencing contact also in that region; if this was not possible then an adjacent white matter electrode was selected; in the case where more than one pair of adjacent channels were eligible, we selected those that had the least amount of epileptic activity according to visual inspection.

Based on the above-mentioned anatomical and functional criteria, one pair of hippocampal depth electrode contacts was selected for each of the ten participants, which were either CA1 or adjacent subfields (**Supplementary Fig. 1**). The number of hippocampal contacts was small in most of our participants, as most of them contained only one pair of hippocampal electrodes. For that reason, and to ensure comparability between regions, we decided to select only one electrode per participant on all cortical areas used in this analysis. As the recordings were conducted stereotactically, we opted to focus on one pair of electrodes in the temporal lobe as well. This choice allowed us to explore neocortical activity patterns and ensured that we could conduct within-participant comparisons in the present study. The decision to include an electrode in the temporal lobe was driven, in part, by the aim to maintain a comparable sampling of regions outside the hippocampus. Additionally, the anterior temporal lobe has been implicated in the processing of visual information during recognition memory (Pacheco Estefan et al. 2019), to represent information about various categories including objects and people (Peelen and Kastner 2011; Tsantani et al. 2019; Reagh and Ranganath 2023) and to interact with hippocampus during successful episodic memory formation (Griffiths et al. 2019). It is important to acknowledge that electrode selection in human studies is constrained by clinical considerations, limiting our choices. Nevertheless, we also added data from an additional pair of electrodes in frontal regions from 6 of the 10 participants, who happened to have an electrode in this region due to clinical considerations.

### Intracranial EEG preprocessing and Ripple Detection

Intracranial analyses were performed to identify ripples and examine their relationship to LFPs. In order to detect ripples, the procedure applied in Vaz et al. 2019; 2020 was used. First, the EEG signal was band-pass filtered in the ripple band (80-120 Hz) using a second order Butterworth filter. Then a Hilbert transformation was applied to extract the instantaneous amplitude within that band. Events were selected if the Hilbert envelope exceeded 2 standard deviations above the mean amplitude of the filtered traces. Only events that were at least 25 ms in duration and had a maximum amplitude greater than 3 standard deviations were retained as ripples for analysis. Adjacent ripples separated by less than 15 ms were merged. Additionally, we ensured that, for each individual candidate, the band-pass signal had at least three peaks and at least three troughs, and that its’ power spectrum exhibited a global peak between 80 Hz and 120 Hz. For this last step the power spectrum was computed for frequencies between 30 Hz and 190 Hz in steps of 2 Hz (using Morlet wavelets with seven cycles) and divided by the power spectrum estimated across the entire recording.

Simultaneously, an automated event-level artifact rejection (Vaz et al. 2019; 2020) was applied to remove system-level line noise, eye-blink artifacts, sharp transients, and interictal epileptiform discharges (IEDs), which can be mistakenly characterized as ripples after high pass filtering. To do so, we calculated a z-score for every time point based on the gradient (first derivative) and amplitude after applying a 250 Hz high pass filter. Any time point that exceeded a z-score of 5 with either gradient or high-frequency amplitude was marked as artifactual, including periods of 200ms before and after each identified time point.

All data and identified ripples were visually inspected to ensure that the above methodology reliably identified ripples and excluded IEDs and apparent high-frequency oscillations associated with IEDs.

For each ripple, we extracted its peak time as the time point at which the band-pass signal was highest; the ripple duration as the time difference between the start and end time of a given ripple; and the inter-ripple interval (IRI) as the time difference between the onset of two consecutive ripples. To depict the time-domain signal, we extracted the raw LFP traces and the time-frequency-domain power spectrum (using Morlet wavelets with 7 cycles at 30 logarithmically spaced frequencies between 1 and 200 Hz), within −100 to 100 ms around each ripple.

### Ripple phase locking to ongoing neural oscillations at the slow band

To investigate whether ripples were locked to particular phases of slow oscillations (0.5 to 4 Hz) we filtered the signal using a two-pass Butterworth filter and extracted the instantaneous phase using the Hilbert transform at the onsets of each ripple. To assess phase consistency across ripples we computed inter-trial phase coherence (ITPC) values across ripples for each participant (Cohen 2014). ITPC spans from 0 to 1, with 1 corresponding to a perfect inter-trial coherence (i.e., the same phase on each trial onset).

To assess the statistical significance of ripple-phase coupling, we compared the empirical values against 1000 surrogate values computed by permuting the inter-ripple interval distribution (i.e., permuting the time differences between the onset of two consecutive ripples), for each participant. Then we computed a group-level p-value of the empirical average ITPC z-value in comparison to the surrogate ITPC z-values as the fraction of surrogate values that were larger than the empirical value, with an alpha of 0.05.

### iEEG spectral power during ripples

To assess whether hippocampal/cortex ripples were associated with significant changes in LFP power in neocortex/hippocampus, respectively, we computed ripple-aligned time frequency-resolved power spectrograms (Kunz et al. 2024) across the entire recording, using Morlet wavelets with 7 cycles at 50 logarithmically spaced frequencies between 1 and 200Hz. Power values were z-scored across time for each frequency. Values around each hippocampal ripple (±3 s) were extracted and time points with IEDs were excluded (i.e., set to NaN). Finally, power z-values were averaged across ripples and smoothed with a Gaussian filter across time (kernel length, 0.2 s). This procedure was computed individually for each participant and then averaged across participants. For visualization, we truncated the spectrogram ±0.5 s around the ripple peak time point.

To statistically evaluate power changes, we performed a cluster-based permutation test (1000 surrogates) across channels in which we first applied a one-sample t-test to the empirical data, separately for each time-frequency bin, and identified contiguous clusters of time-frequency bins in which the uncorrected p-value of the t-test was significant (α = 0.05). Then for each cluster, we computed an empirical cluster statistic by summing up all t-values being part of that cluster. The empirical cluster statistics was compared against surrogate cluster statistics, obtained by flipping the sign of the power values of a random subset of the spectrograms, performing the same steps as for the empirical data, and keeping only the maximum cluster (Kunz et al. 2024). The empirical cluster statistic was considered significant if it exceeded the 95th percentile or if it fell below the 5th percentile of all surrogate maximum cluster statistics.

### Ripple rate during the encoding of movie events

The analysis of the ripple rate during the encoding of movie events was assessed by counting the number of ripples that occurred within each event, for each participant. This value was normalized by the length of the event, and then the resulting normalized ripple count was averaged across events. To evaluate the extent to which the number of ripples during the encoding of an event determined its successful recall at the later verbal recall test, for each participant, we split the events that were later recalled and forgotten and obtained an averaged measure of ripple count for each condition and participant, which was then compared by using a one-sample paired t-test, with significance threshold set at an alpha of 0.05.

### Ripple rate at movie event boundaries

To assess how ripples fluctuated around event boundaries we computed a peri-ripple time histogram of ripples across event boundaries, for each participant. We used 300-ms time bins starting from –2 to 2 s relative to boundary occurrence. For visualization purposes, we smoothed it with a 5-point triangular window. This resulted in an estimated probability of observing a ripple at each time-point during the −2 to 2 s epoch.

The empirical ripple rates were compared against 1000 surrogate values computed by calculating peri-stimuli histograms of the −2 to 2 s epoch but by shuffling the temporal order of the events throughout the movie. This procedure ensured that each surrogate preserved signal properties and maintained the lengths of the events, resulting in ripple rates that corresponded to within event windows, for each participant. For each time point surrounding an event boundary, we determined the p-value by calculating the fraction of surrogate values that exceeded the empirical value, considering the z-value of the empirical rate. An alpha level of 0.05 was used for this analysis. The p-values were later FDR corrected (Benjamini and Hochberg 1995).

## Notes

### Competing Interest Statement

The authors have declared no competing interest.

### Summary of Updates

This version of the manuscript reflects the results after some extra control measures were added to the ripple detection algorithm.

