## SupplementaryFigures for "Movie-watching evokes ripple-like activity within events and at event boundaries"

### Supplementary Figures

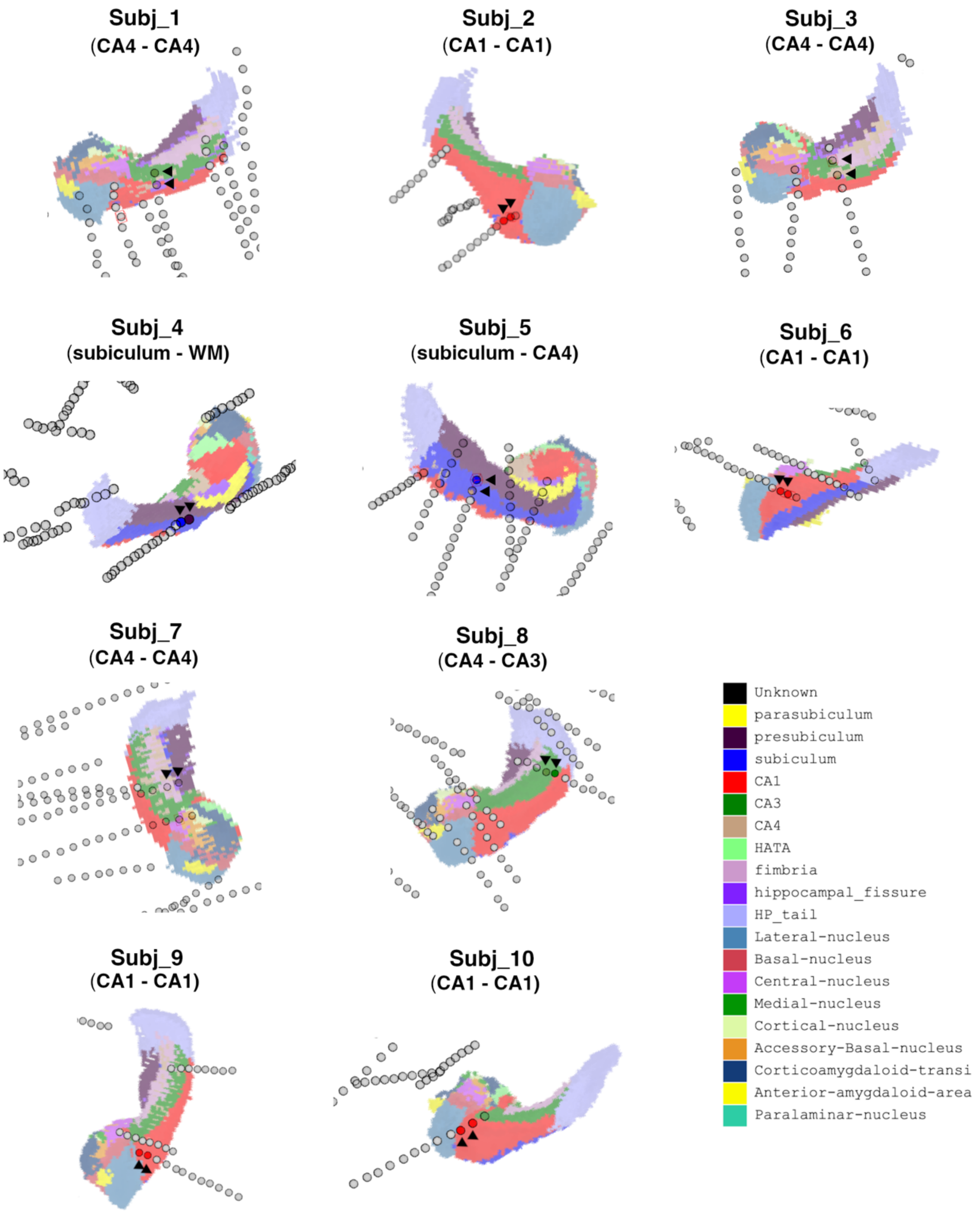


**Supplementary Figure 1:** Depiction of the hippocampal macroelectrode channels for each participant. Hippocampus is devided in its different subregions. Electrode channels used for the analysis are colored with the respective subregion they were implated in and indicated by black arrows. Other electrode channels not used in the study can be seen in grey.

**Table 1:** Subjects information and SOZ.

| | Age | Gender | Electrode Hemisphere | SOZ |
|---|---|---|---|---|
| **Subject 1** | 43 | F | Left | Left insula |
| **Subject 2** | 18 | M | Right | Posterior quadrant and left temporal lobe dysplasia areas |
| **Subject 3** | 28 | M | Left | Right occipital dysplasia |
| **Subject 4** | 22 | M | Left | Temporal pole |
| **Subject 5** | 24 | F | Left | Left posterior insula |
| **Subject 6** | 30 | M | Left | Left mesial frontal |
| **Subject 7** | 43 | F | Right | Temporal pole |
| **Subject 8** | 30 | M | Left | Left superior frontal peri-encephalomalacia |
| **Subject 9** | 33 | M | Left | Left anterior-mesial frontal |
| **Subject 10** | 20 | F | Left | Anterior dysplasia involving left anterior and media insula |

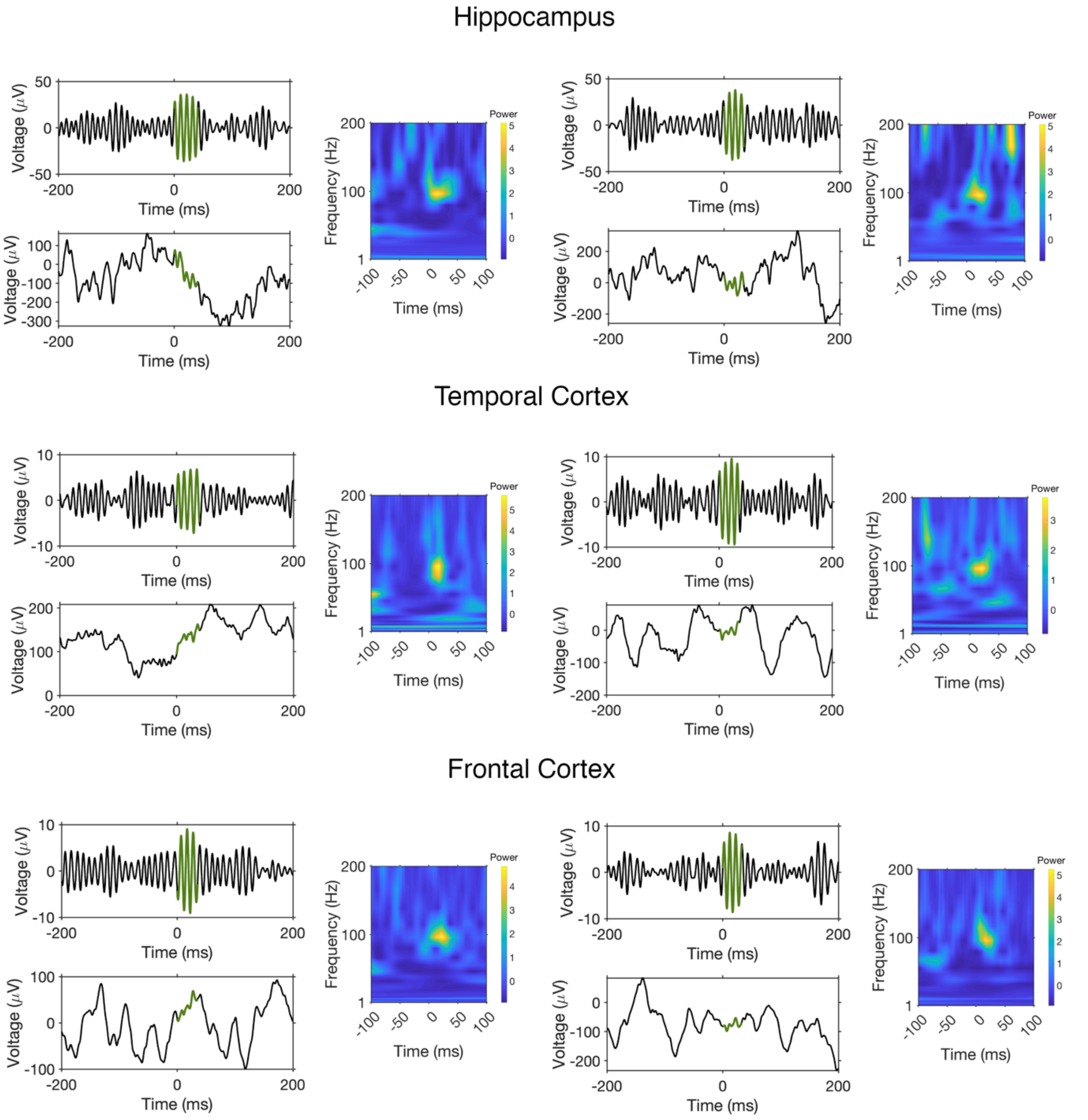


**Supplementary Figure 2:** Raw LFP, LFP filtered in the 80–120 Hz ripple band and z-scored power spectrogram in the time-frequency domain for some example ripples.

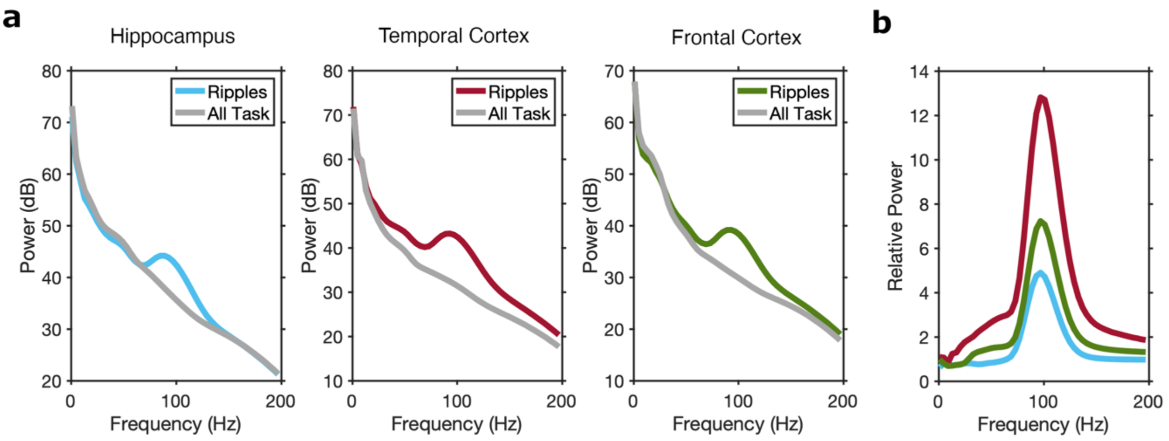


**Supplementary Figure 3: (a)** Average power spectrum of ripples (colored) and average power spectrum for the all task (grey). **(b)** Relative power estimated by dividing the power values during the ripples by the mean power values across the entire experiment (separately for each frequency). As expected, ripple periods showed peaks at around 90 Hz.

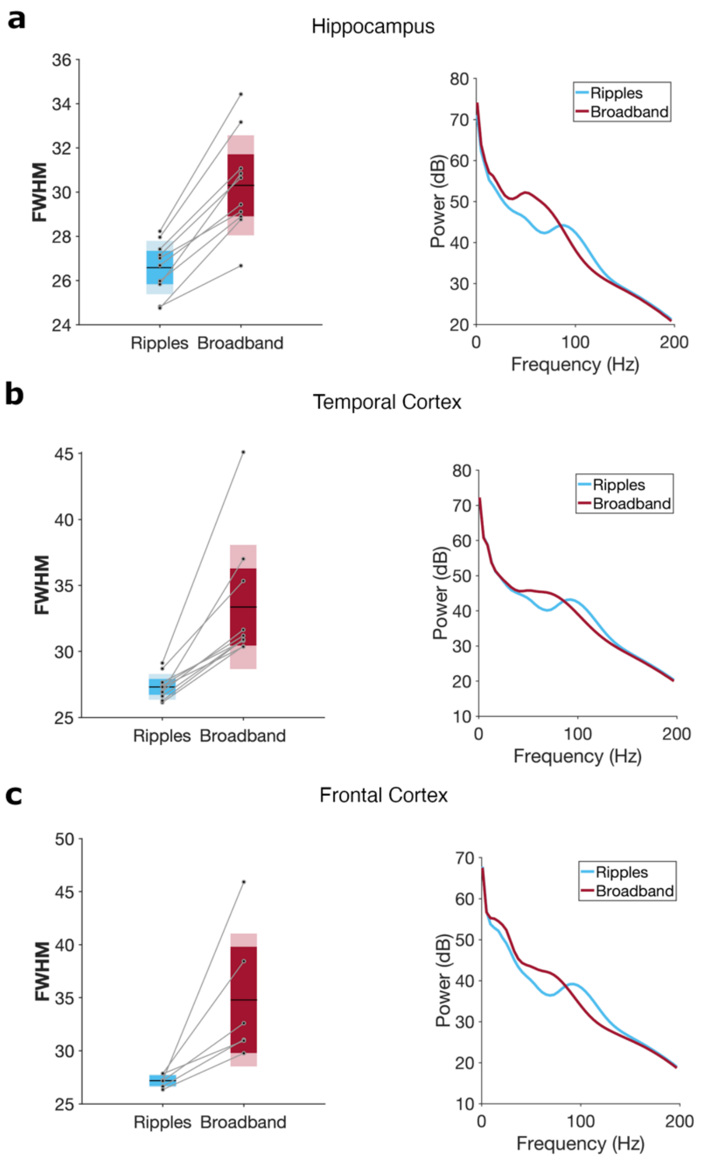


**Supplementary Figure 4:** Frequency spread for ripples and broadband activity events (left), and average power spectrum of these events (right) for **(a)** hippocampus, **(b)** temporal cortex and **(c)** frontal cortex. Broadband events were computed by running the ripple algorithm with a larger bandpass (50 - 180Hz) and a lower threshold of event acceptance (1SD instead of the 2SD used for ripple detection). For all these events (ripples and broadband) we have: (1) extracted the time period of each event; (2) computed the time-frequency spectrum of the data during this period using Morlet wavelets with 7 cycles between 1 and 200Hz; (3) averaged across time to obtain a frequency spectrum for each event; (4) computed the Full width at half maximum (FWHM) for the spectrum above 30Hz**.** The FWHM of ripples was significantly lower than the broadband events (Rank-sum Test; $p < 0.001$, for all regions). Additionally, these results could not be explained by decreased frequency resolution during time-frequency decomposition with wavelets as the broadband events occur at a lower frequency than ripples.

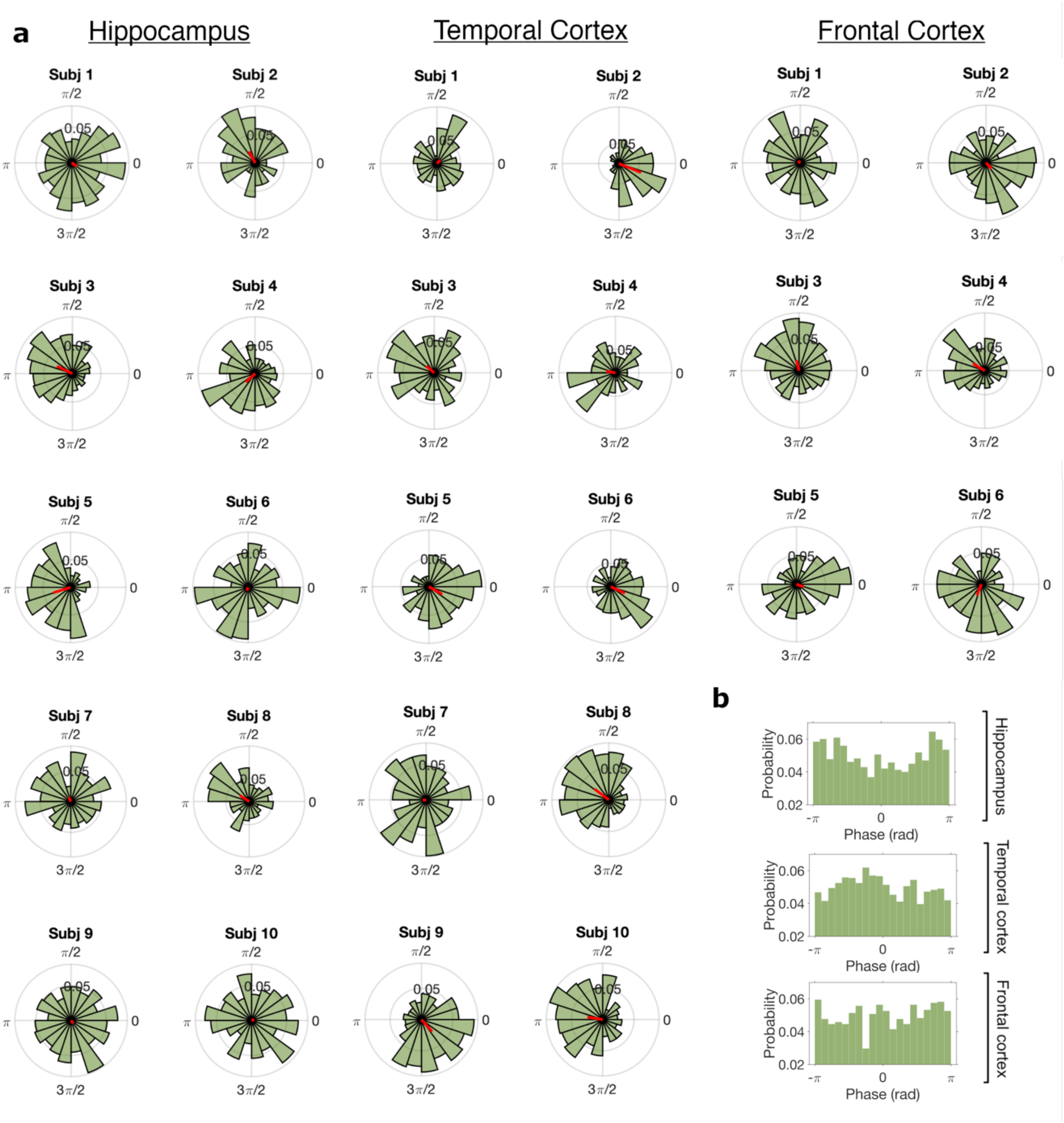


**Supplementary Figure 5:** Ripple low oscillations locking phase for individual participants. **a)** Polar distribution of phase angles at the onset of the hippocampal, temporal cortex and frontal cortex ripples, for each participant. Grand average across ripples is depicted by the thick red line. **b)** Histogram distribution of slow oscillation phase angles at the onset of the hippocampal, temporal cortex and frontal cortex ripples, for all participants.

**Table 2:** Results of Linear Mixed Model where Model 1: Random intercept only, Model 2: Random intercept + Slope (Region), Model 3: Random intercept + Slope (Memory), Model 4: Random intercept + Slopes (Region + Memory). Models 3 and 4 indicated with an asterisk, exhibited a singular fit. The selected model is highlighted in bold.

| *Model* | *Contrast* | *Df numerator* | *Df denominator* | *F* | *p* |
|---|---|---|---|---|---|
| 1 | MemoryTag | 1 | 963.460 | 1.244 | .265 |
| 1 | Region | 2 | 958.531 | 4.709 | .009 |
| 1 | MemoryTag * Region | 2 | 956.199 | 3.683 | .026 |
| **2** | **MemoryTag** | **1** | **915.777** | **.933** | **.334** |
| **2** | **Region** | **2** | **7.109** | **.508** | **.622** |
| **2** | **MemoryTag * Region** | **2** | **922.628** | **3.253** | **.039** |
| 3* | MemoryTag | 1 | 22.660 | .993 | .330 |
| 3* | Region | 2 | 939.791 | 4.698 | .009 |
| 3* | MemoryTag * Region | 2 | 947.174 | 3.821 | .022 |
| 4* | MemoryTag | 1 | 23.641 | .728 | .402 |
| 4* | Region | 2 | 7.251 | .518 | .616 |
| 4* | MemoryTag * Region | 2 | 866.030 | 3.267 | .039 |

**Table 3:** Results from the Bayesian Information Criterion (BIC) analysis performed to compare the different Linear Mixed Models that varied in the inclusion of random slopes for each fixed effects (none, Memory only, Region only, both). The selected model is highlighted in bold.

| *Model* | *BIC* |
|---|---|
| Data ~ Memory * Region+ (1\|Subject) | -1664.510 |
| **Data ~ Memory * Region+ (1+Region\|Subject)** | **-1688.332** |

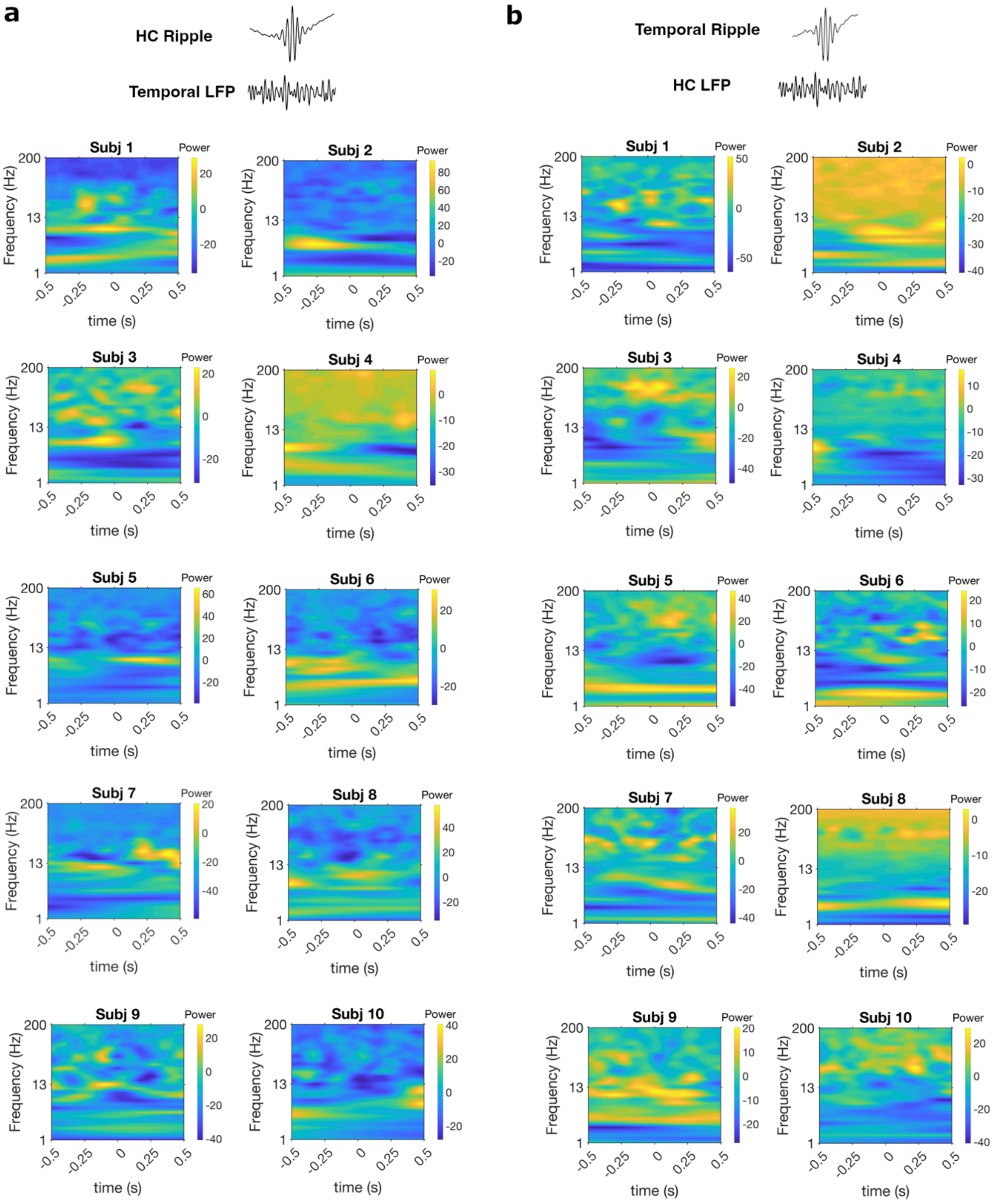


**Supplementary Figure 6:** (**a**) LFP power (z-scored) changes in temporal cortex locked to hippocampal ripples for each participant. (**b**) LFP power (z-scored) changes in hippocampus locked to temporal cortex ripples for each participant. Data in all above analysis was smoothed with a gaussian filter across time (kernel length, 0.2 s) and 0 corresponds to ripple peak.

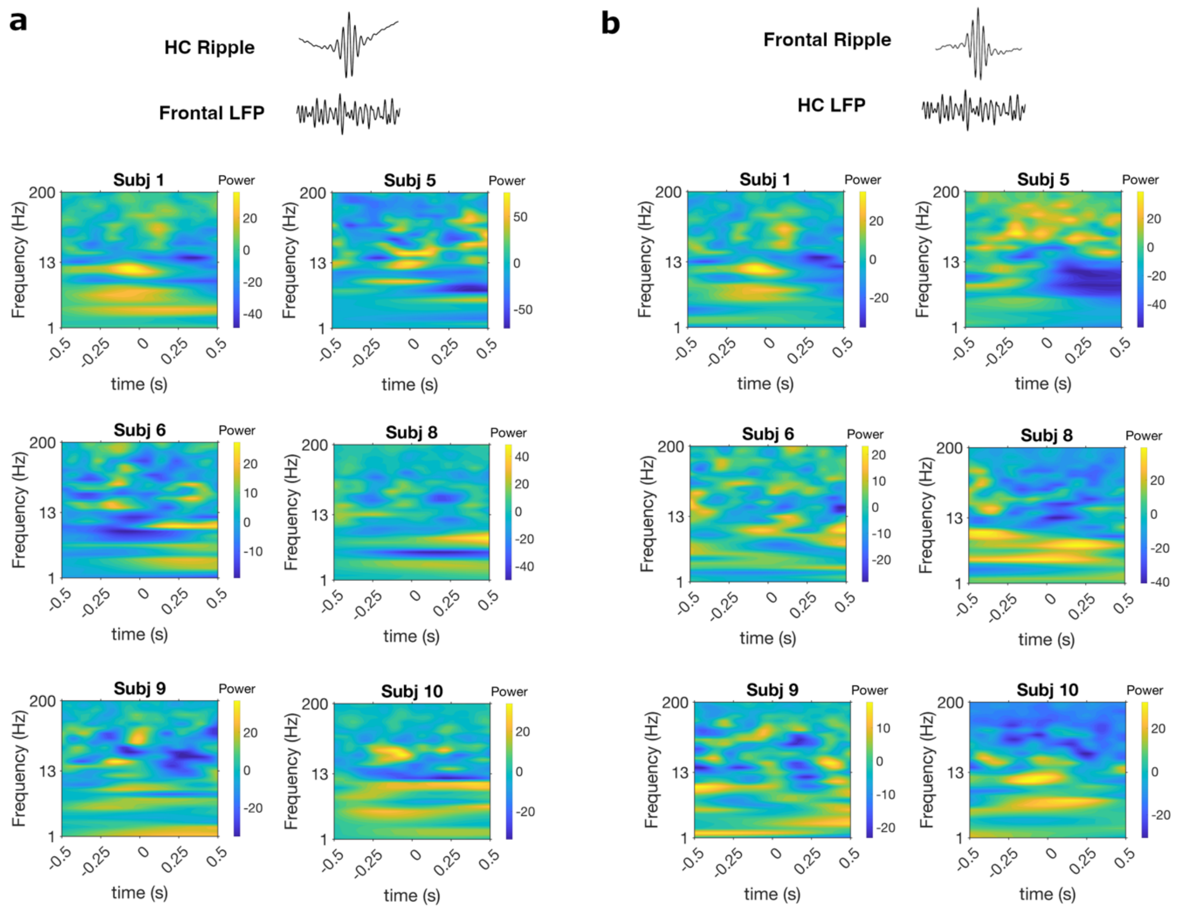


**Supplementary Figure 7:** (**a**) LFP power (z-scored) changes in frontal cortex locked to hippocampal ripples for each participant. (**b**) LFP power (z-scored) changes in hippocampus locked to frontal cortex ripples for each participant. Data in all above analysis was smoothed with a gaussian filter across time (kernel length, 0.2 s) and 0 corresponds to ripple peak.

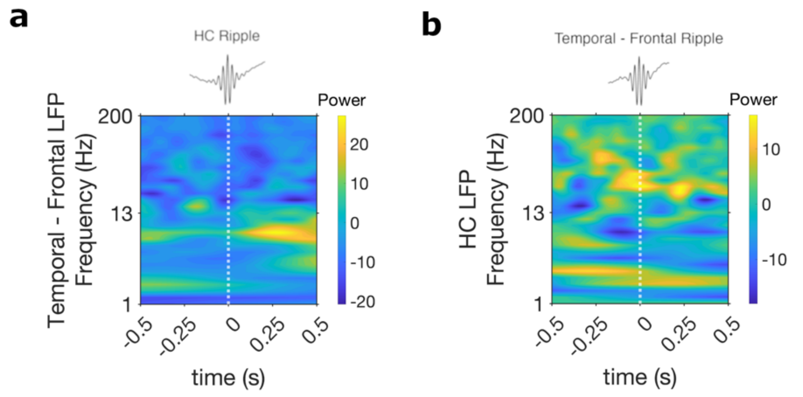


**Supplementary Figure 8:** LFP power (z-scored) difference **(a)** between temporal cortex and frontal cortex LFP timed to hippocampal ripple occurrence and **(b)** hippocampal LFP changes between moments of temporal cortex ripple occurrence and frontal cortex ripple occurrence. Data in all above analysis was smoothed with a gaussian filter across time (kernel length, 0.2 s).

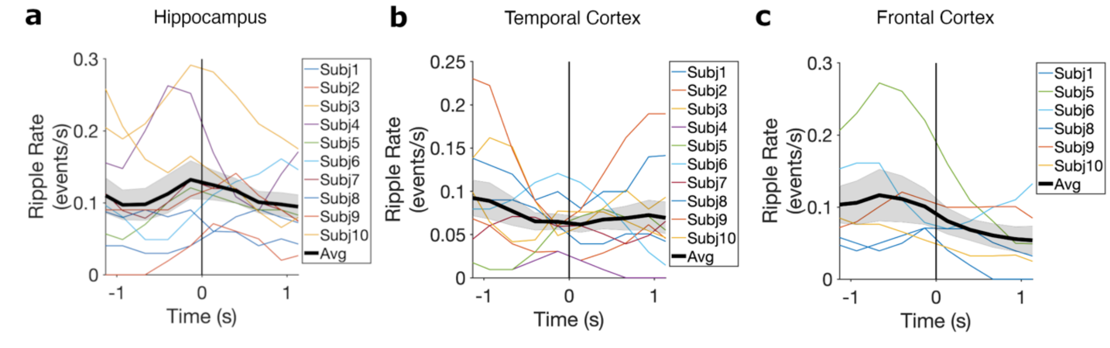


**Supplementary Figure 9:** Instantaneous ripple rate computed in 300ms time bins around boundary onset and smoothed by a five-point triangular window, for each participant data (colored) and average data (black), for **a)** hippocampus, **b)** temporal cortex and **c)** frontal cortex. Shaded region corresponds to SEM across participants.

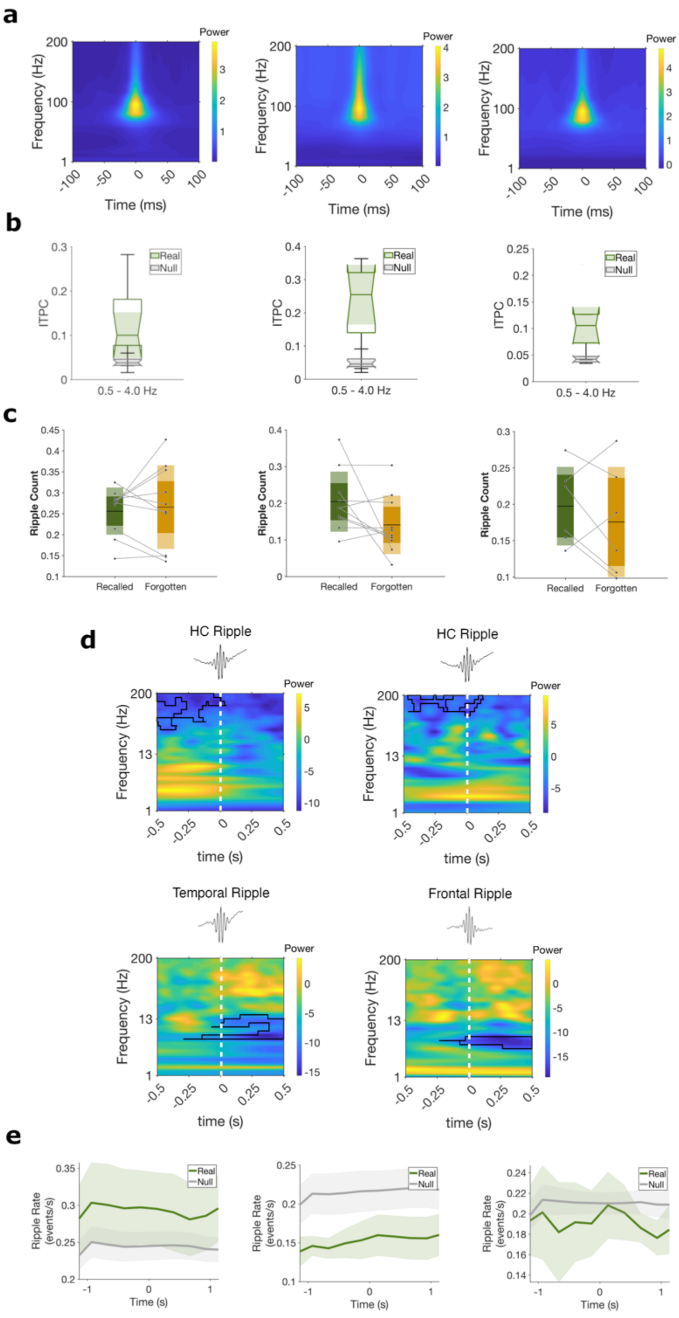


**Supplementary Figure 10:** Results after applying a different ripple detection method. (**a**) Grand-average z-scored power spectrogram in the time-frequency domain, with time 0 corresponding to ripple peak, for (left) hippocampus, (center) temporal cortex and (right) frontal cortex. (**b**) Inter-trial phase coherence (ITPC) values across ripples (green) and surrogate data (gray), for (left) hippocampus, (center) temporal cortex and (right) frontal cortex. **(c)** Average frequency of ripples during an event, for each participant, normalized by the length of the event, for recalled (green) and forgotten (yellow) events, in hippocampus (left), temporal cortex (center) and frontal cortex (right) For all boxplots, the central mark is the median, and the edges of the box are the 25th and 75th percentiles. **(d)** LFP power (z-scored) changes in (top-left) temporal cortex and (top-right) frontal cortex locked to hippocampal ripples, where 0 corresponds to ripple peak. LFP power (z-scored) changes in hippocampus locked to (bottom-left) temporal cortex and (bottom-right) frontal cortex ripples during the encoding of the movie, where 0 corresponds to ripple peak. **(e)** Instantaneous ripple rate computed in 300ms time bins around boundary onset and smoothed by a five-point triangular window, for empirical data (green) and surrogate data (grey), for (left) hippocampus, (center) temporal cortex and (right) frontal cortex. Shaded region corresponds to SEM across participants.
